# Identifying the cortical face network with dynamic face stimuli: A large group fMRI study

**DOI:** 10.1101/2023.09.26.559583

**Authors:** David Pitcher, Geena R Ianni, Kelsey Holiday, Leslie G Ungerleider

**Author notes:** Corresponding author: David Pitcher -, Department of Psychology, University of York, Heslington, York, YO10 5DD, U.K.

## Abstract

Functional magnetic resonance imaging (fMRI) studies have identified a network of face-selective regions distributed across the human brain. In the present study, we analyzed data from a large group of gender-balanced participants to investigate how reliably these face-selective regions could be identified across both cerebral hemispheres. Participants (*N*=52) were scanned with fMRI while viewing short videos of faces, bodies, and objects. Results revealed that five face-selective regions: the fusiform face area (FFA), posterior superior temporal sulcus (pSTS), anterior superior temporal sulcus (aSTS), inferior frontal gyrus (IFG) and the amygdala were all larger in the right than in the left hemisphere. The occipital face area (OFA) was larger in the right hemisphere as well, but the difference between the hemispheres was not significant. The neural response to moving faces was also greater in face-selective regions in the right than in the left hemisphere. An additional analysis revealed that the pSTS and IFG were significantly larger in the right hemisphere compared to other face-selective regions. This pattern of results demonstrates that moving faces are preferentially processed in the right hemisphere and that the pSTS and IFG appear to be the strongest drivers of this laterality. An analysis of gender revealed that face-selective regions were typically larger in females (*N*=26) than males (*N*=26), but this gender difference was not statistically significant.

## Introduction

Functional magnetic resonance imaging (fMRI) studies of face processing reliably identify multiple face-selective regions distributed across the human brain (Gauthier et al., 2000; Kanwisher et al., 1997; McCarthy et al., 1997; Puce et al., 1996; Puce et al., 1998). Each of these face-selective regions is thought to preferentially process a different facial aspect. For example, the fusiform face area (FFA) preferentially processes individual identity (Grill-Spector et al., 2004), the posterior superior temporal sulcus (pSTS) preferentially processes emotional expressions (Winston et al., 2004) and the occipital face area (OFA) preferentially processes the parts of a face, such as the eyes and mouth (Gauthier et al., 2000; Pitcher et al., 2007). Cognitive and neurobiological models have proposed that these regions form the components of a distributed network specialized for processing faces (Bruce & Young, 1986; Calder & Young, 2005; Haxby et al., 2000). In the present study, we quantified the extent to which face-selective regions could be reliably identified with fMRI across both cerebral hemispheres in a large group of experimental participants (*N*=52) using moving face stimuli.

Previous fMRI studies have demonstrated that moving face stimuli increase the reliability of localizing face-selective regions, particularly the pSTS (Fox et al., 2009; LaBar et al., 2003; Schultz & Pilz, 2009), the anterior STS (aSTS) and the inferior frontal gyrus (IFG) (Pitcher et al., 2011). This has led to proposals that the STS contains cortical regions that preferentially process changeable aspects of other people’s faces, such as their emotional state (Allison et al., 2000) and their attentional focus as revealed by their gaze direction (Calder et al., 2007). However, the extent to which face-selective regions are lateralized to the right hemisphere when participants view moving faces is still unclear. A recent study using moving faces showed that the pSTS was more right lateralized in humans than in macaques (De Winter et al., 2015) but the extent to which the amygdala and the inferior frontal gyrus (IFG) are also lateralized in humans is still unclear. Evidence from different experimental methodologies have demonstrated that faces are preferentially processed in the right hemisphere (Barton et al., 2002; Kanwisher et al., 1997; Pitcher et al., 2007) but all of these studies used static faces as stimuli. Given the strong preference for moving faces in the STS, we used videos of faces to localize face-selective regions, in order to compare STS regions with other face-selective regions in the human brain. Our aim was to investigate how reliably each face-selective region can be identified across both the right and left hemispheres.

Participants were scanned, using fMRI, while they viewed short videos of faces, bodies, and objects. We then examined whether face-selective regions (defined using a contrast of activation by faces greater than objects) in the fusiform gyrus, the inferior occipital gyrus, the posterior and anterior superior temporal sulcus, the amygdala, and the inferior frontal gyrus were present in both hemispheres. To further characterize these regions we also measured the neural response in these regions-of-interest (using independent data) to these same face videos, as well as moving bodies and objects (as control stimuli). The participant group was gender balanced (females *N*=26, males *N*=26) so we additionally looked to see whether there were any gender differences in the reliability and size of face-selective regions. Our results demonstrated that the FFA, pSTS, aSTS, amygdala, and IFG were all significantly larger in the right than the left hemisphere; the OFA showed the same pattern but the difference failed to reach significance. However, the right hemispheric advantage for moving faces was larger in the pSTS and IFG than in other face-selective regions. Consistent with prior evidence, we also observed that face-selective regions were larger and more likely to be present in females, but these differences were not statistically significant.

## Materials and Methods

### Participants

A total of 52 right-handed participants (26 females, 26 males) with normal, or corrected-to-normal, vision gave informed consent as directed by the National Institutes of Health (NIH) Institutional Review Board (IRB).

### Stimuli

#### Regions-of-interest (ROIs) Localizer Stimuli

Face-selective regions-of-interest (ROIs) were identified using 3-second video clips from three different stimulus categories (faces, bodies, and objects). These videos had been used in previous fMRI studies of face perception (Pitcher et al., 2011; Pitcher et al., 2019; Sliwinska, Bearpark, et al., 2020; Sliwinska, Elson, et al., 2020). There were sixty video clips for each category in which distinct exemplars appeared multiple times. Videos of faces and bodies were filmed on a black background and framed close-up to reveal only the faces or bodies of 7 children as they danced or played with toys or with adults (who were out of frame). Fifteen different moving objects were selected that minimized any suggestion of animacy of the object itself or of a hidden actor moving the object (these included mobiles, windup toys, toy planes and tractors, balls rolling down sloped inclines). Stimuli were presented in categorical blocks and, within each block, stimuli were randomly selected from the entire set for that stimulus category. This meant that the same actor or object could appear within the same block. Participants were instructed to press a button when the subject in the stimulus was repeated in the same block (i.e. a repeat of the same actor, body, or object). The order of repeats was randomized and happened an average of once per block.

### Procedure

Functional data were acquired over 6 blocked-design functional runs lasting 234 seconds each. Each run contained three sets of three consecutive stimulus blocks (faces, bodies, and objects) alternating with rest blocks, to make three blocks per stimulus category per run. Each block lasted 18 seconds and contained stimuli from one of the three stimulus categories. Stimulus order was pseudo-randomized such that each category appeared once every three blocks but, within each of these blocks of three, the stimulus category order was randomized. Each functional run contained four 18-second rest blocks, one each at the beginning and end of the run, one at 72 seconds and another at 144 seconds. During rest blocks, a series of six uniform colored fields were presented for three seconds each. Participants were asked to perform a 1-back working memory task in which they were required to press a response button when the subject of the video occurred twice in a row. After functional data were acquired we also collected a high resolution T-1 weighted anatomical scan to localize the functional activations.

### Brain Imaging and Analysis

Participants were scanned on a research dedicated GE 3-Tesla scanner. Whole brain images were acquired using a 32-channel head coil (36 slices, 3 × 3 × 3 mm, 0.6 mm interslice gap, TR = 2 s, TE = 30 ms). Slices were aligned with the anterior/posterior commissure. In addition, a high-resolution T-1 weighted MPRAGE anatomical scan (T1-weighted FLASH, 1 x 1 x 1 mm resolution) was acquired to anatomically localize functional activations.

Functional MRI data were analyzed using AFNI (http://afni.nimh.nih.gov/afni). Data from the first four TRs from each run were discarded. The remaining images were slice-time corrected and realigned to the third volume of the first functional run and to the corresponding anatomical scan. The volume-registered data were spatially smoothed with a 5-mm full-width-half-maximum Gaussian kernel. Signal intensity was normalized to the mean signal value within each run and multiplied by 100 so that the data represented percent signal change from the mean signal value before analysis.

A general linear model (GLM) was established by convolving the standard hemodynamic response function with the 3 regressors of interest (one for each stimulus category - faces, bodies, and objects). Regressors of no interest (e.g., 6 head movement parameters obtained during volume registration and AFNI’s baseline estimates) were also included in this GLM.

We initially performed a group whole brain analysis using 3 runs each from all participants (the same three runs used for the ROI analysis, namely runs 2, 4 and 6). Data were entered into a random-effects ANOVA with faces, bodies, and objects as fixed factors and participants as a random factor. Statistical maps were calculated at a threshold of *p* =0.001 and a cluster correction of fifty voxels. Face-selective regions were identified using a contrast of activation by faces greater than objects.

We then performed an ROI analysis to further characterize the data. The six functional runs were divided in two for analysis purposes. Even runs (runs 2, 4 and 6) were used to identify face-selective ROIs, odd runs (runs 1, 3 and 5) were used to calculate the magnitude or response to faces, bodies and objects in these ROIs. Face-selective ROIs were identified for each participant using a contrast of activation by faces greater than objects, calculating significance maps of the brain using a statistical threshold of *p* = 0.0001. Within each functionally defined ROI, we then calculated the magnitude of response (percent signal change from a fixation baseline) for faces, bodies, and objects.

## Results

### Group whole brain analysis

We first performed a whole brain group analysis to illustrate the extent of face-selective cortex across both hemispheres. Data from all participants (*N* = 52) was entered into a random-effects ANOVA with the three stimulus categories (faces, bodies, and objects) as fixed factors and participants as a random factor. Statistical maps were calculated at a threshold of *p* =0.0001 and a cluster correction of fifty voxels. The results from a contrast of activation by faces greater than objects are shown in Figure 1. We observed more face-selective voxels in the right hemisphere than in the left hemisphere. This was most striking along the superior temporal sulcus where there was no corresponding significant activation in the left hemisphere.

**Figure 1.**
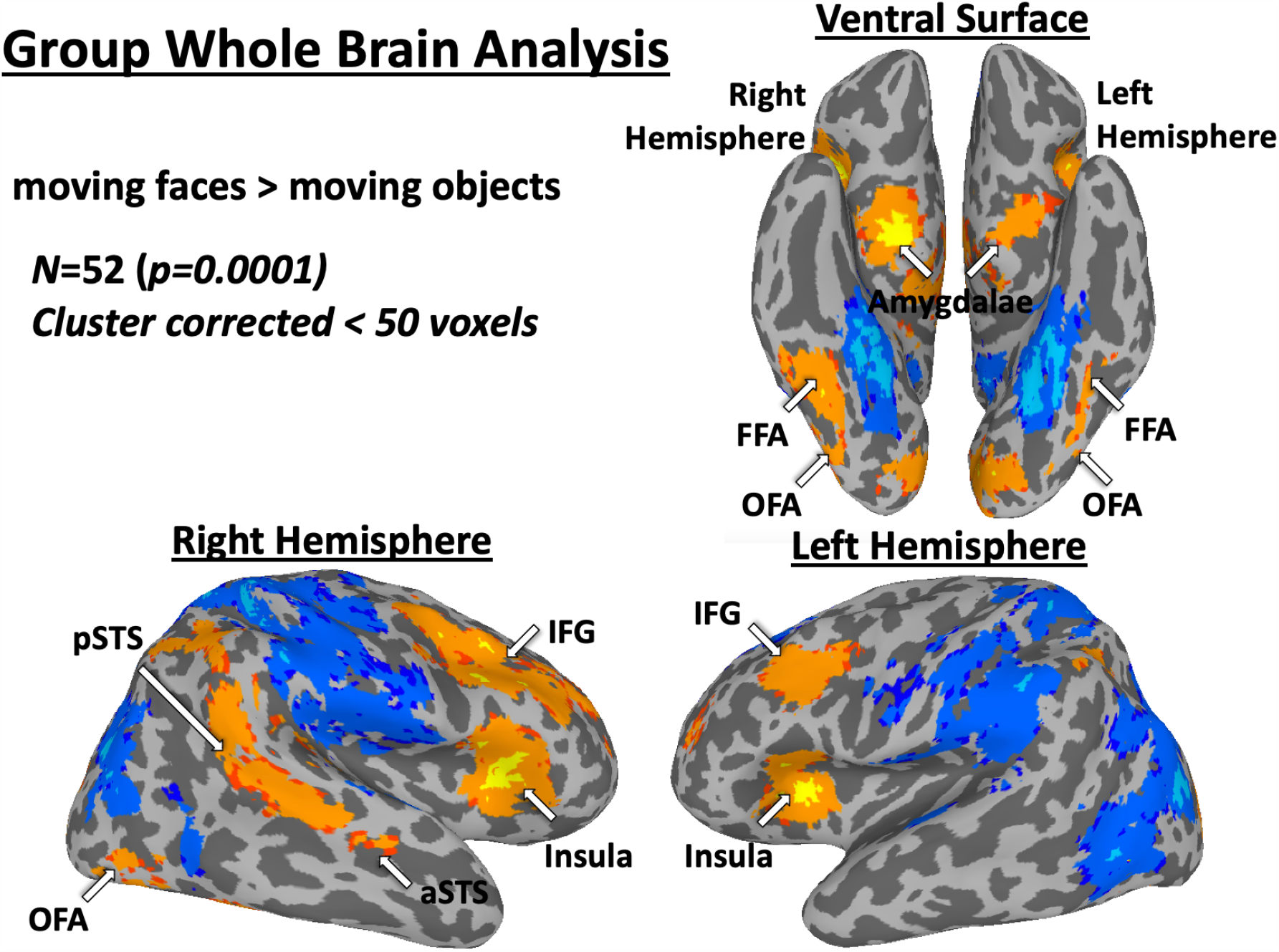
A whole-brain group analysis (*N*=52) showing a contrast of activation by faces greater than objects.

#### ROI analysis

Face-selective ROIs were identified based on the data from three runs of the localizer using a contrast of activation by faces greater than objects. Consistent with prior literature, face-selective ROIs in the left hemisphere (LH) were less common and smaller across participants than in the right hemisphere (Kanwisher et al., 1997; Barton et al., 2002; Young et al., 1986; Pitcher et al., 2007; 2011). The number of participants exhibiting face-selective ROIs across both hemispheres together with the mean MNI co-ordinates of each ROI is shown in Table 1.

**Table 1.**
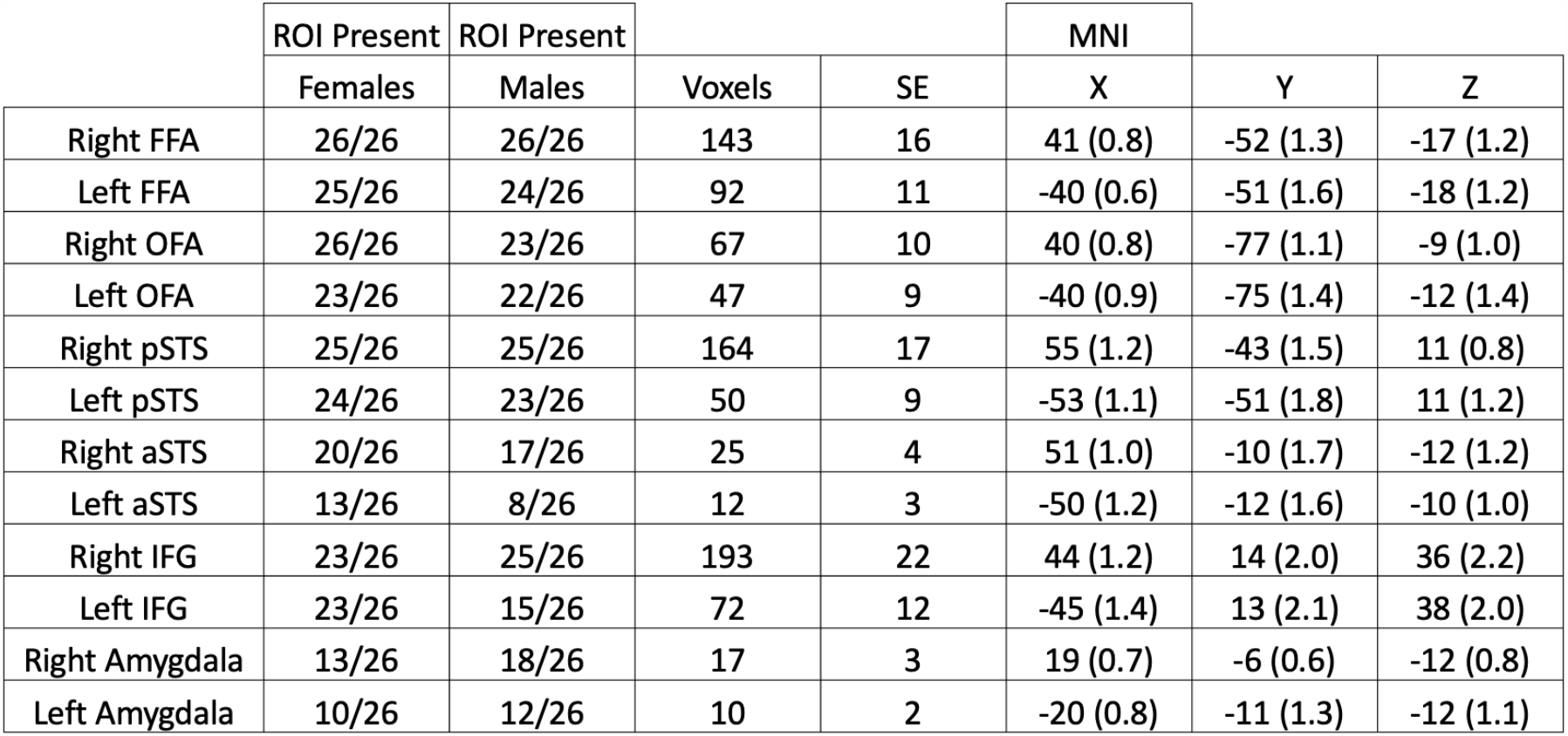
Results of the ROI analysis. Consistent with prior evidence, face-selective ROIs were identified more consistently and were larger in size in the right than in the left hemisphere.

To quantify the size of face-selective ROIs across the two hemispheres, we measured the number of voxels in each of the face-selective ROIs (Table 1, Figure 2). These data were entered into a two (hemisphere: right, left) by six (ROI: FFA, OFA, pSTS, aSTS, amygdala, and IFG) repeated measures ANOVA. Results showed significant main effects of hemisphere (F (1,51)=71, *p* **<** 0.0001) and ROI (F (5,255)=32, *p* **<** 0.0001) and there was a significant interaction between the two factors (F (5,255)=18, *p* **<** 0.0001). Post-hoc tests showed that there were more voxels in the right than in the left hemisphere in the FFA (*p* **=** 0.001), the pSTS (*p* **<** 0.0001), the aSTS (*p* **=** 0.009), the amygdala (*p* **=** 0.001) and the IFG (*p* **<** 0.0001). A similar trend was found in the OFA, but the difference did not reach significance (*p* **=** 0.065).

**Figure 2.**
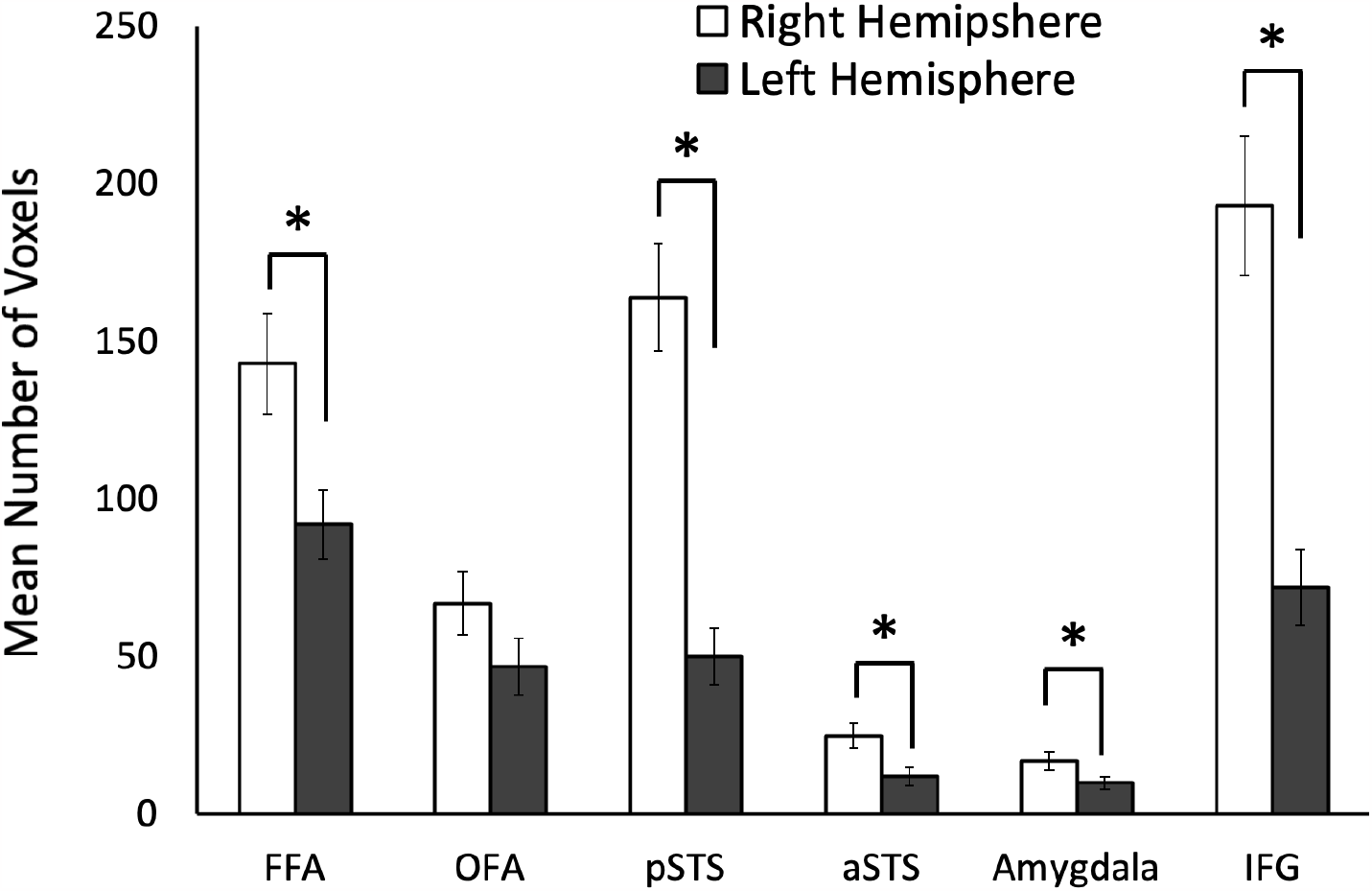
The mean size of the six face-selective ROIs, as measured by number of voxels calculated from all fifty-two participants. Error bars denote across participants standard error of the mean (*SE*). Results showed that all face-selective ROIs were significantly larger in the right than the left hemisphere, with the exception of the OFA, which did not reach significance (*p*=0.065).

To further quantify the extent of the hemispheric laterality effects across face-selective ROIs, we calculated a laterality index. To do this we subtracted the number of voxels in the left hemisphere ROI from the number of voxels from the corresponding right hemisphere ROI in each participant (see Figure 3). We then entered these data into a one-way ANOVA, which revealed a significant main effect of ROI (F (5,255)=18, *p* **<** 0.0001). Post-hoc comparisons showed that the pSTS and IFG were significantly more right lateralized than all other ROIs (*p* < 0.04) and that the FFA was significantly more right lateralized than the aSTS (*p* = 0.016) and amygdala (*p* = 0.039).

**Figure 3.**
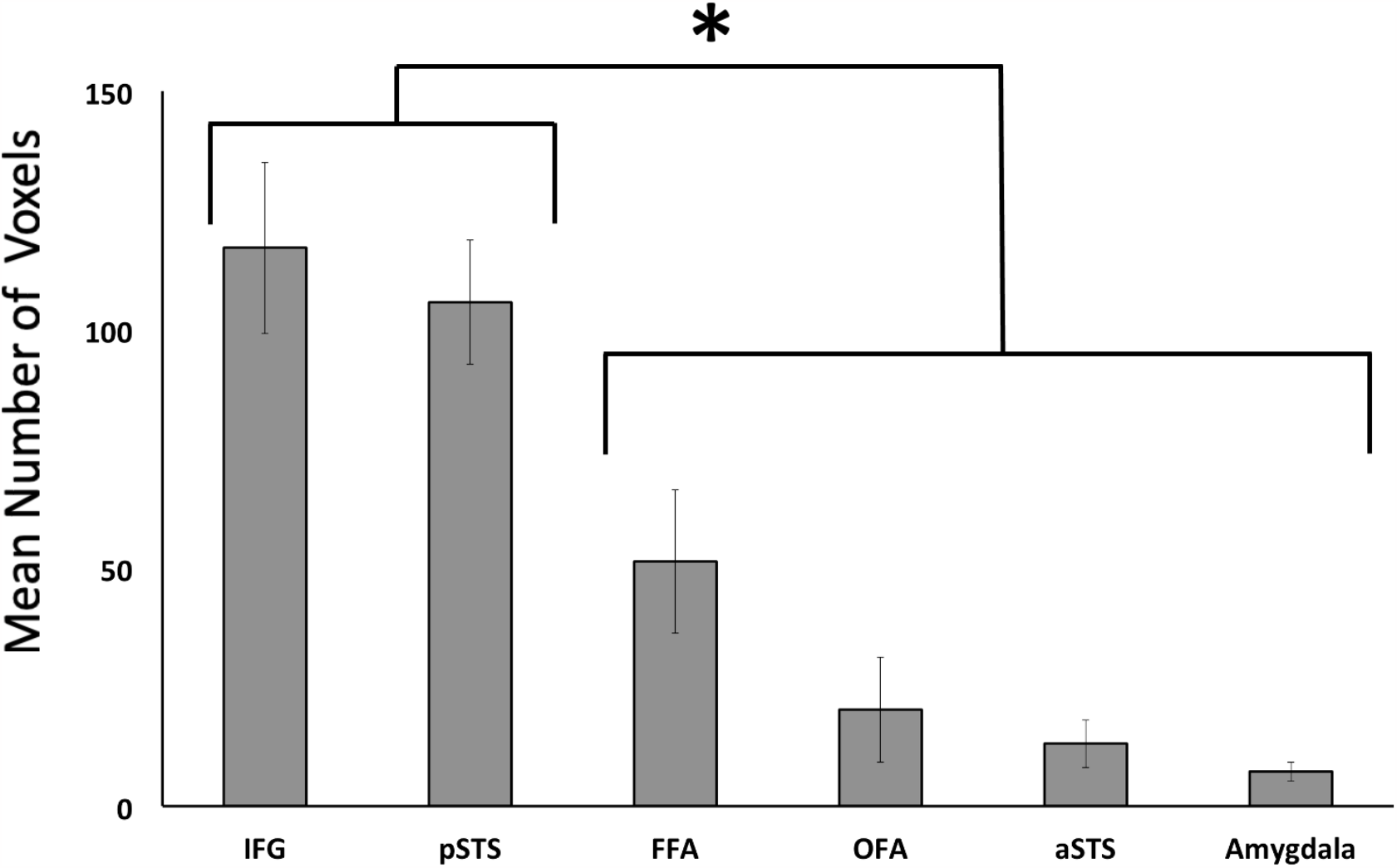
The right laterality index of the six face-selective ROIs. This was calculated by subtracting the size of the left hemisphere ROI from the corresponding ROI in the right hemisphere. Results showed that the IFG and pSTS were significantly larger in the right hemisphere than the four other face-selective ROIs. Error bars denote across participant standard error of the mean (*SE*).

#### ROI Response Profiles to faces, bodies and objects

We next examined the response profiles of each of the face-selective ROIs (i.e., rFFA, rOFA, rpSTS, raSTS, amygdala, and IFG) to short videos of faces, bodies, and objects using the independently calculated percent signal change data (see Figure 4). The percent signal change data was entered into a two (hemisphere: right, left) by three (stimulus category: faces, bodies, and objects) by six (ROI: FFA, OFA, pSTS, aSTS, amygdala, and IFG) repeated measures ANOVA. Results showed significant main effects of stimulus category (F (2,24)=24, *p* **<** 0.0001) and ROI (F (5,60)=23, *p* **<** 0.0001) but not of hemisphere (F (1,12)=1.4, *p* = 0.3). There were also interactions between hemisphere and stimulus category (F (2,24)=3.2, *p* **=** 0.035) and between ROI and stimulus category (F (10,120)=15, *p* **<** 0.0001). These interactions revealed that faces elicited a greater response in the right than in the left hemisphere ROIs and that faces exhibited a greater response than bodies and objects across all ROIs. No further interactions approached significance (*p* > 0.7).

**Figure 4.**
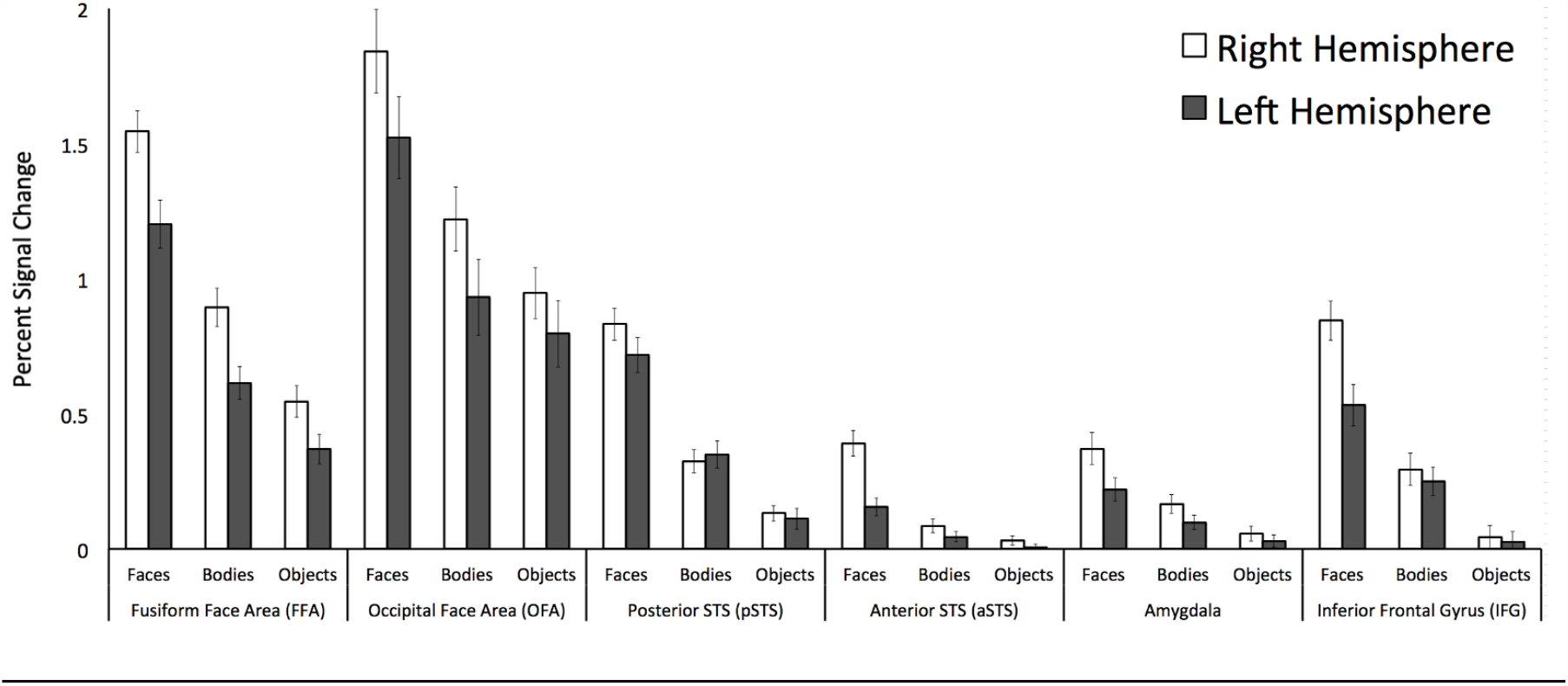
Percent signal change data for faces, bodies and objects in the face-selective ROIs in both hemispheres. Results showed a significant main effect of stimulus and that faces produced a significantly greater response in the right than in the left hemisphere. Error bars denote across participant standard error of the mean (*SE*).

#### Gender Analysis

To quantify the size of face-selective ROIs across the two genders, we measured the number of voxels in each of the face-selective ROIs of the female (*N*=26) and male participants (*N*=26) (Figure 5). These data were entered into a two (gender: female, male) by two (hemisphere: right, left) by six (ROI: FFA, OFA, pSTS, aSTS, amygdala, and IFG) repeated measures analysis of variance (ANOVA). Results showed significant main effects of hemisphere (F (1,25)=105, *p* **<** 0.0001) and ROI (F (5,125)=29, *p* **<** 0.0001) but not of gender (F (1,25)=2.2, *p* **=** 0.14). There was also a significant interaction between hemisphere and ROI (F (5,125)=18, *p* **<** 0.0001) showing that face-selective ROIs were larger in the right than in the left hemisphere. No other interactions approached significance (*p* **>** 0.3).

**Figure 5.**
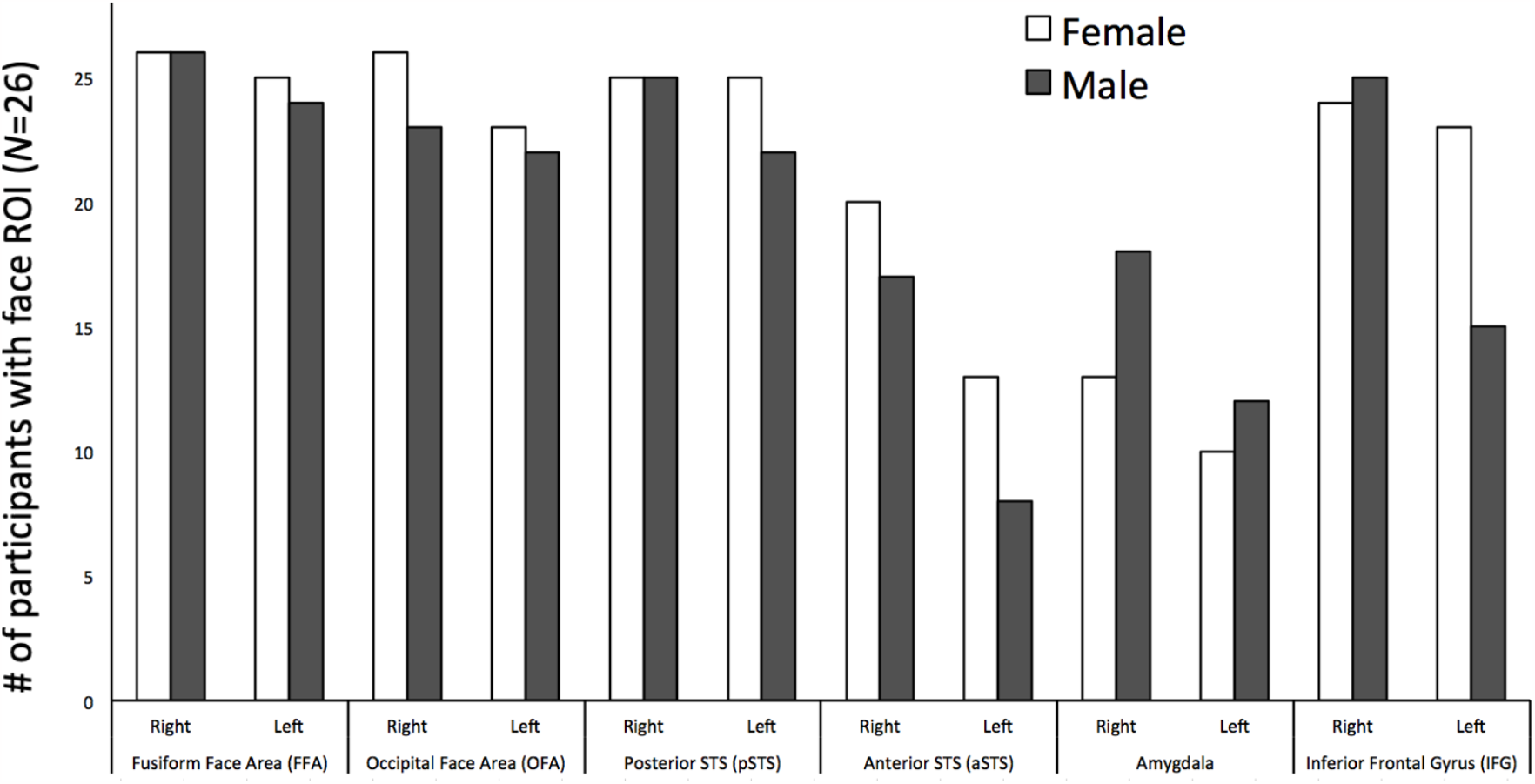
The mean size of the six face-selective ROIs as measured by number of voxels calculated separately for female (*N*=26) and male (*N*=26) participants. Results showed a significant main effect of ROI and of hemisphere but not of gender, and a significant interaction between ROI and hemisphere. No other interactions approached significance (error bars denote across subject standard error of the mean).

## Discussion

The results of our large group fMRI study clearly demonstrate that dynamic, moving faces are preferentially processed in the right hemisphere. Face-selective regions were larger in the right than the left hemisphere and, additionally, the neural response to faces was greater in the right than the left hemisphere ROIs. This pattern of results is consistent with prior studies that used static face images as stimuli (Barton et al., 2002; Bona et al., 2015; Kanwisher et al., 1997; Landis et al., 1986; Pitcher et al., 2007; Young et al., 1986; Yovel et al., 2003). Intriguingly, we observed a greater right-hemisphere laterality in the pSTS and IFG than in the other face-selective regions. This suggests that regions that exhibit a stronger response to dynamic faces (Allison et al., 2000; Nikel et al., 2022; Pitcher et al., 2011; Pitcher & Ungerleider, 2021) show a greater degree of hemispheric specialization than regions that show little, or no, preference for dynamic over static faces, such as the FFA and OFA (Fox et al., 2009; Pitcher et al., 2011; Pitcher et al., 2019; Schultz & Pilz, 2009; Sliwinska, Bearpark, et al., 2020). This greater degree of right lateralization in the pSTS and IFG (Figure 3) is also strikingly demonstrated by our group whole brain analysis (Figure 1). We observed a large and contiguous face-selective activation in the right hemisphere running from the parietal lobe along entire STS; by contrast, no face-selective voxels were observed in the left STS.

Neuroimaging studies have shown that the pSTS and aSTS exhibit a greater response to dynamic than static faces (Puce et al., 1998; LaBar et al., 2002; Schulz et al., 2012) while the FFA and OFA show little, or no preference for dynamic over static faces (Fox et al., 2009; Pitcher et al., 2011). Our results demonstrate that the pSTS is also more right lateralized than the FFA and OFA. It is unclear why a face-selective region that preferentially processes moving over static faces should be more lateralized but one suggestion may come from the functional role of the left STS. Many studies have demonstrated that cortical regions near the left STS, such as the left middle temporal gyrus, are preferentially engaged by language-related tasks (Vigneau et al., 2006). A recent study demonstrated that face-selectivity in the posterior STS was more right lateralized in humans than in macaques, suggesting that language processing may have led to hemispheric specialization in humans (De Winter et al., 2015). In addition, we demonstrated that TMS disrupts expression discrimination in the right pSTS more than in the left pSTS (but this study used static face stimuli) (Sliwinska & Pitcher, 2018). Interestingly, we observed that the left pSTS was located more posteriorly than the right pSTS (Table 1), which seems consistent with language-selective cortex being more prominent than face-selective cortex in left temporal brain regions.

We also observed that the face-selective ROI in the IFG was significantly larger in the right than the left hemisphere. The functional role of the IFG in the face network is not currently clear, but a prior study demonstrated that it responded more strongly to moving than static faces like the pSTS (Pitcher et al., 2011). Consistent with this result we have further demonstrated that both the IFG and pSTS exhibit on equal response to faces in both visual fields while the OFA and FFA exhibit a contralateral bias (Nikel et al., 2022; Pitcher et al., 2020). The IFG has been implicated in face working memory studies (Courtney et al., 1996), famous face recognition (Ishai, 2002) and in preferentially processing information from the eyes (Chan & Downing, 2011). The greater degree of laterality we observed in the right IFG and pSTS is interesting and suggests that the two regions may be functionally connected to perform similar cognitive functions and warrants further investigation. For example, a recent large-scale analysis of data from 680 participants reported that the IFG and STS formed a face sub-network specialized for the processing of dynamic facial information.

Prior neuroimaging studies have suggested that men show a greater degree of hemispheric specialization for language (Shaywitz et al., 1995) and faces (Proverbio et al., 2006; Wager et al., 2003) compared to women. It is also worth noting that prior divided visual field behavioural studies reported that both genders exhibit a right hemisphere advantage for face processing tasks, but the laterality of this effect is larger in men than women (Bourne, 2005; Young & Bion, 1983). Our participant group was gender balanced (26 females, 26 males) so, in addition to the whole group analysis, we also examined whether there were any gender differences in size and reliability of face-selective ROIs. While we observed larger face-selective ROIs in females than males (Figure 5), this difference was not statistically significant. We also observed that females had more face-selective ROIs in the pSTS and aSTS, while men were more likely to have face-selective voxels in the amygdala (Table 2), but these differences were also not statistically significant. It may be that our sample was too small to detect small but reliable gender differences and future work may be required.

In the present study, we presented data collected from a large group of participants who were scanned with fMRI while viewing moving face, body, and object stimuli. Results demonstrated that the moving faces are preferentially processed in the right hemisphere; this was demonstrated both in the size of face-selective ROIs and in the neural response to moving faces. Additional analysis revealed that this right lateralization for dynamic faces was stronger in the pSTS and IFG than in other face-selective regions. Our results suggest that the functional role of the IFG in the face network warrants further investigation.

## Acknowledgements

The research reported here was supported by the Intramural Research Program of the NIMH (NCT01087281; 93-M-0170 and NCT01617408; 12-M-0128) and by a grant from the Simons Foundation to D.P. (SFARI# 392150). We thank Nancy Kanwisher for providing stimuli and Philip Quinlan for statistical advice.

